# Heading Perception Depends on Time-Varying Evolution of Optic Flow

**DOI:** 10.1101/356758

**Authors:** Charlie S. Burlingham, David J. Heeger

## Abstract

There is considerable support for the hypothesis that perception of heading in the presence of rotation is mediated by instantaneous optic flow. This hypothesis, however, has never been tested. We introduce a novel method, termed “non-varying phase motion,” for generating a stimulus that conveys a single instantaneous optic flow field, even though the stimulus is presented for an extended period of time. In this experiment, observers viewed stimulus videos and performed a forced choice heading discrimination task. For non-varying phase motion, observers made large errors in heading judgments. This suggests that instantaneous optic flow is insufficient for heading perception in the presence of rotation. These errors were mostly eliminated when the velocity of phase motion was varied over time to convey the evolving sequence of optic flow fields corresponding to a particular heading. This demonstrates that heading perception in the presence of rotation relies on the time-varying evolution of optic flow. We hypothesize that the visual system accurately computes heading, despite rotation, based on optic acceleration, the temporal derivative of optic flow.

---

James Gibson first remarked that the instantaneous motion of points on the retina (Fig. 1a) can be formally described as a two-dimensional field of velocity vectors called the “optic flow field” (or “optic flow”) (1). Such optic flow, caused by an observer’s movement relative to the environment, conveys information about self-motion and the structure of the visual scene (1–15). When an observer translates in a given direction along a straight path, the optic flow field radiates from a point in the image with zero velocity, or singularity, called the focus of expansion (FOE; Fig. 1b). It is well known that under such conditions, one can accurately estimate one’s instantaneous direction of translation or “heading” by simply locating the FOE (see *SI Appendix*). However, if there is angular rotation in addition to translation (by moving along a curved path or by a head or eye movement), the singularity in the optic flow field will be displaced such that it no longer corresponds to the true heading (Fig. 1c, d). In this case, if one estimates heading by locating the singularity, the estimate will be biased away from the true heading. This is known as the rotation problem (14).

**Fig. 1.**
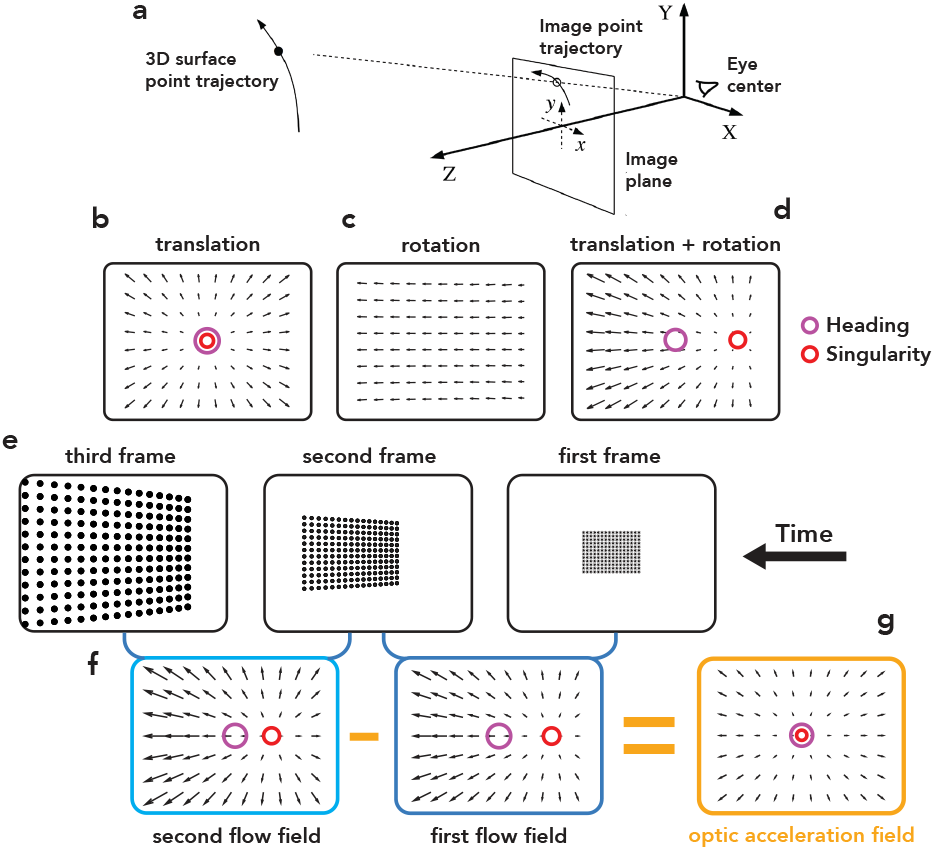
Projective geometry, the rotation problem, time-varying optic flow, and the optic acceleration hypothesis. **a.** Viewer-centered coordinate frame and perspective projection. Owing to motion between the viewpoint and the scene, a 3-D surface point traverses a path in 3-D space. Under perspective projection, the 3-D path of this point projects onto a 2-D path in the image plane (retina), the temporal derivative of which is called image velocity. The 2-D velocities associated with all visible points defines a dense 2-D vector field called the optic flow field. **b, c, d.** Illustration of the rotation problem. **b.** Optic flow for pure translation (1.5 m/s translation speed, 0° heading, i.e., heading in the direction of gaze). Optic flow singularity (red circle) corresponds to heading (purple circle). **c.** Pure rotation, for illustrative purposes only and not corresponding to any experimental condition (2 °/s rightward rotation). **d.** Translation + rotation (1.5 m/s translation speed, 0° heading, 2 °/s rightward rotation). Optic flow singularity (red circle) is displaced away from heading (purple circle). **e.** Three frames from a video depicting movement along a circular path with the line-of-sight initially perpendicular to a single fronto-parallel plane composed of black dots. **f.** Time-varying evolution of optic flow. The first optic flow field reflects image motion between the first and second frames of the video. The second optic flow field reflects image motion between the second and third frames of the video. For this special case (circular path), the optic flow field evolves (and the optic flow singularity drifts) only due to the changing depth of the environment relative to the viewpoint. **g.** Illustration of the optic acceleration hypothesis. Optic acceleration is the derivative of optic flow over time (here, approximated as the difference between the second and first optic flow fields). The singularity of the optic acceleration field corresponds to the heading direction. Acceleration vectors scaled by 10x for visibility.

Computer vision researchers and vision scientists have developed a variety of algorithms that accurately and precisely extract observer translation and rotation from optic flow, thereby solving the rotation problem. Nearly all of these rely on the instantaneous optic flow field (4, 9, 16–25) with few exceptions (26–28). However, it is unknown whether these algorithms are commensurate with the neural computations underlying heading perception.

The consensus of opinion in the experimental literature is that human observers can estimate heading (29, 30) from the instantaneous optic flow field, in the absence of additional information (5, 10, 15, 31–33). Even so, there are reports of systematic biases in heading perception (11); the visual consequences of rotation (eye, head, and body) can bias heading judgments (10, 15, 34–36), with the amount of bias typically proportional to the magnitude of rotation. Other visual factors such as stereo cues (37, 38), depth structure (8, 10, 39–42), and field of view (32, 41–43) can modulate the strength of these biases. Errors in heading judgments have been reported to be greater when eye (34–36, 44, 45) or head movements (36) are simulated versus when they are real, which has been taken to mean that observers *require* extra-retinal information, although there is also evidence to the contrary (10, 15, 32, 39, 40, 43, 46–49). Regardless, to date, no one has tested whether heading perception (even with these biases) is based on instantaneous optic flow or on the information available in how the optic flow field evolves over time. Some have suggested that heading estimates rely on information accumulated over time (31, 43, 50), but no one has investigated the role of time-varying optic flow without confounding it with stimulus duration (i.e., the duration of evidence accumulation).

In this study, we employed a novel application of an image processing technique that ensured that only a single instantaneous optic flow field was available to observers, even though the stimulus was presented for an extended period of time. We called this condition “non-varying phase motion” or “non-varying”: the phases of two component gratings comprising each stationary stimulus patch shifted over time at a constant rate, causing a percept of motion in the absence of veridical movement (51). Phase motion also eliminated other cues that may otherwise have been used for heading judgments including image point trajectories (15, 31) and their spatial compositions (i.e., looming (52, 53) and dynamic perspective distortions (46, 54)). For non-varying phase motion, observers exhibited large biases in heading judgments, in the presence of rotation. A second condition, “time-varying phase motion” or “time-varying,” included acceleration by varying the velocity of phase motion over time to match the evolution of a sequence of optic flow fields. Doing so allowed observers to compensate for the confounding effect of rotation on optic flow, making heading perception nearly veridical. This demonstrates that heading perception in the presence of rotation relies on the time-varying evolution of optic flow.

## Results

Observers (N = 10) travelled on a virtual circular path, following previous studies (10, 15, 42), and performed a forced-choice heading discrimination task. They judged whether heading was left or right of the center of the field of view, and were provided “correct”/“incorrect” feedback after each trial. Heading (relative to center) was constant over time within each trial. Stimuli conveyed headings from −20° (left) to +20° (right), where 0° was center, and simultaneous rotations of −2, −0.8, 0 (no-rotation), +0.8, or +2 °/s angular velocity, corresponding to circular paths of different radii. The virtual environment was initialized as two fronto-parallel planes positioned at 12.5 and 25 m, following a previous study (10). Heading bias was computed from the center of the psychometric function (50:50 leftward:rightward choices) and compared across conditions (Fig. 3). Stimulus videos contained varying levels of information about the time evolution of the optic flow field (Fig. 2). There were three stimulus conditions: non-varying phase motion, time-varying phase motion, and envelope motion. Non-varying phase motion conveyed only instantaneous optic flow — a single optic flow field that was either the first or last in an evolving sequence — but for an extended duration (see *Movies S1-4*). Time-varying phase motion conveyed time-varying optic flow — the entire evolving sequence of optic flow fields (Fig. 1f, *Movies S5-6*). Neither non-varying nor time-varying phase motion showed image point trajectories. Envelope motion (i.e., “dot motion”) conveyed three cues: instantaneous optic flow, time-varying optic flow, and image point trajectories (see *Movies S7-8*).

**Fig. 2.**
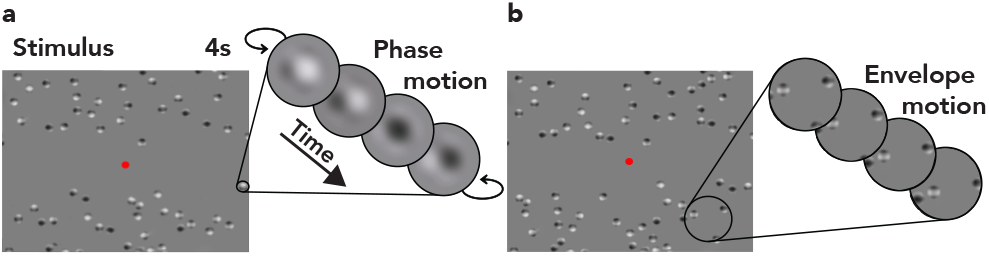
Stimulus conditions. Each panel displays a single frame of a stimulus video from that condition, with an inset (magnified in scale) depicting how a small region of the stimulus changed over time. **a.** Phase motion. For non-varying phase motion, the envelopes of the plaid patches were fixed in location, but the phases of each component within each patch shifted with a constant rate. For time-varying phase motion, the phases shifted with a time-varying rate (i.e., the phase motion accelerated/decelerated over time). **b.** Envelope motion. The envelopes of the plaid patches moved over time, but the phases within them were fixed. For illustrative purposes, in both panels, plaid patches have been enlarged by a factor of four, the fixation dot has been enlarged by a factor of six, and dot density has been reduced by a factor of four.

### Heading perception was strongly biased for non-varying phase motion

Heading bias was significantly larger for non-varying phase motion than envelope motion at all non-zero rotation velocities. This was true for both first and last flow field non-varying phase motion (Fig. 4a, Table S1). Judgments of heading were biased in the direction of rotation and the magnitude of bias scaled with the speed of rotation. For ±2 °/s rotations, when pooled across observers, heading bias (absolute, pooled for +/− rotations) was 2.7x larger on average for non-varying phase motion (average over first and last flow field conditions) than for envelope motion (5.2° vs. 1.9°). For the slower rotation speed of 0.8 °/s, bias was ~5.3x larger for non-varying phase motion than for envelope motion (1.6° vs. 0.3°; see Fig. 4a for pooled data and *SI Appendix*, Fig. S1, for each observer’s data). Heading judgments were also more variable for non-varying phase motion than envelope motion (“variability” = 1/slope of psychometric function). For ±2 °/s rotations, they were 1.6x as variable on average (5.9° vs. 3.62°, non-varying vs. envelope; see *SI Appendix*, Fig. S2). These results demonstrate that observers could not perceive heading veridically from instantaneous optic flow (when rotation was present).

### Heading perception was nearly veridical when the time-varying evolution of optic flow was available

Time-varying phase motion controlled for all cues except time-varying optic flow. Although identical in static appearance to non-varying phase motion, the speed of each component grating varied over time to convey an evolving sequence of optic flow fields — the same sequence presented with envelope motion (see Fig. 1f and *Methods*). Thus, envelope motion and time-varying phase motion conveyed common time-varying optic flow. Heading biases for the two were statistically indistinguishable for 2 °/s rotations, when pooled across observers (Fig. 4a; *SI Appendix*, Table S3). Furthermore, heading bias was very low for time-varying phase motion — indistinguishable from veridical (0° heading) for ±0.8 °/s rotations (Fig. 4a). It follows that heading judgments were also significantly less biased for time-varying phase motion than for non-varying phase motion (Fig. 4a; *SI Appendix*, Fig. S1, Table S2), bolstering our claim that instantaneous optic flow is insufficient. Biases for ±2 °/s rotations were on average 3.3x smaller (1.59° vs. 5.2°, time-varying vs. non-varying phase motion). Biases for ±0.8 °/s rotation were on average 10.8x smaller (0.15° vs. 1.62°, time-varying vs. non-varying phase motion). Taken together, these results suggest that heading perception in the presence of rotation depends on time-varying optic flow. Similar biases for envelope motion and time-varying phase motion suggest, consistent with previous reports (15, 31, 38, 55, 56), that image point trajectories (and their spatial compositions) are not a critical cue for heading perception.

The small biases we observed for time-varying stimuli (envelope motion and time-varying phase motion) are consistent in magnitude with those reported in prior studies, for the (relatively slow) rotation speeds that we used (10, 15). In accord with these previous reports, we observed that heading bias scaled with rotation speed (Fig. 4a). Although observers can compensate for the confounding effects of slow speed rotations (< 1 °/s), some better than others (*SI Appendix*, Fig. S1), they often exhibit larger errors for faster rotation speeds (10, 15).

### Heading judgments were better than an optic flow singularity-based strategy, for non-varying phase motion

We created a null model, or “worst case” model of performance (10), that estimates heading as the singularity of optic flow, a biased strategy (Fig. 1b–d; *Methods*), and compared its predictions with observed heading biases (Fig. 3 & 4b). The null model predicted heading bias correctly for a subset of observers (O3, O8, O9, O10) and a subset of rotation velocities, i.e., null model predictions fell within the 95% credible intervals of the observed biases (*SI Appendix*, Fig. S3). This suggests that O3, O8, O9, & O10 may have sometimes estimated heading by simply locating the singularity of optic flow. O3 had large biases for all stimuli (including time-varying), further evidence that O3 may have used this strategy. However, considering the data set as a whole (pooled across observers), heading judgments were somewhat tolerant to rotation. Biases for non-varying phase motion (last) fell halfway between the null model and veridical. Biases for non-varying phase motion (first) fell one-half to one-third of the way between the null model and veridical.

**Fig. 3.**
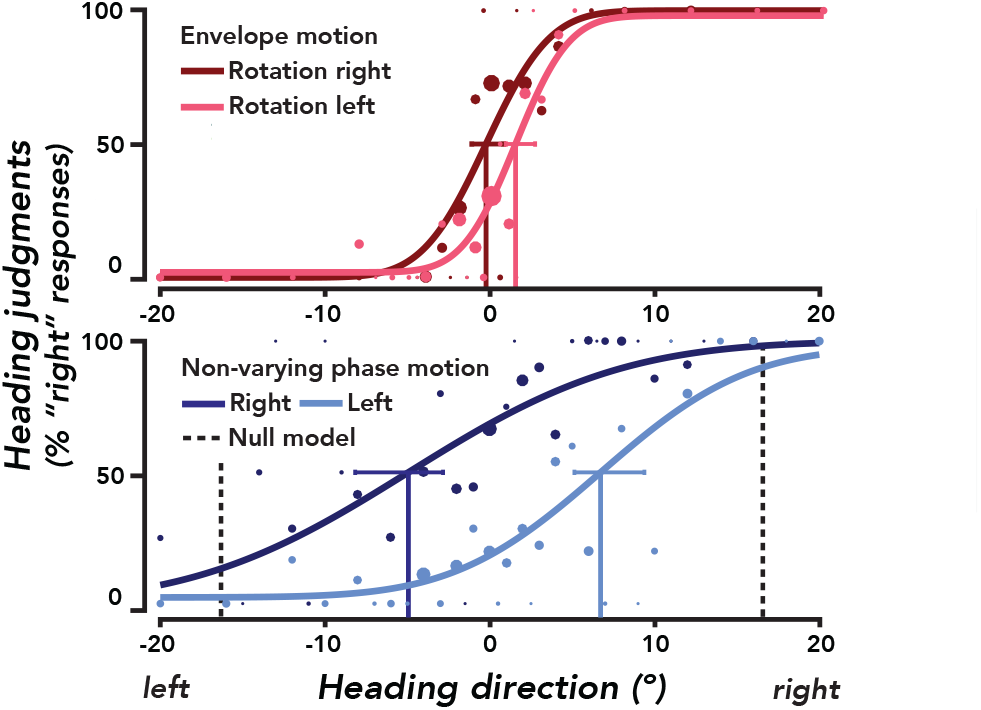
Example psychometric functions for envelope and non-varying phase motion. Data from a single observer (N trials per curve = 240). Data points represent percentage of “right” responses at each heading. Dot area indicates the number of data points collected at that heading. Smooth curves, best-fit psychometric functions. Solid vertical lines indicate bias. Error bars, 95% credible intervals for the bias estimates. Red, envelope motion. Blue, non-varying phase motion. Dark colors, −2 °/s (leftward) rotation. Light colors, +2 °/s (rightward) rotation. Dashed vertical lines, null model predictions.

Heading bias was largely indistinguishable for the first and last optic flow fields, suggesting that each instantaneous optic flow field was equally uninformative. Envelope motion and time-varying phase motion stimuli conveyed a common sequence of optic flow fields. To generate non-varying phase motion stimuli, we picked a single optic flow field from this sequence, either the first or last. On trials with rotation, the singularity of optic flow drifted toward the true heading over time due to the changing depth and angle of the fronto-parallel planes. If observers used an optic flow singularity-based strategy, biases would be larger for non-varying (first) than for non-varying (last). Pooled across observers, heading bias was indistinguishable between the two (Fig. 4b and *SI Appendix*, Table S5; p > 0.04) except for rightward 2 °/s rotation, for which bias was higher for the first than last optic flow field, an effect driven just by O2 and O6.

**Fig. 4.**
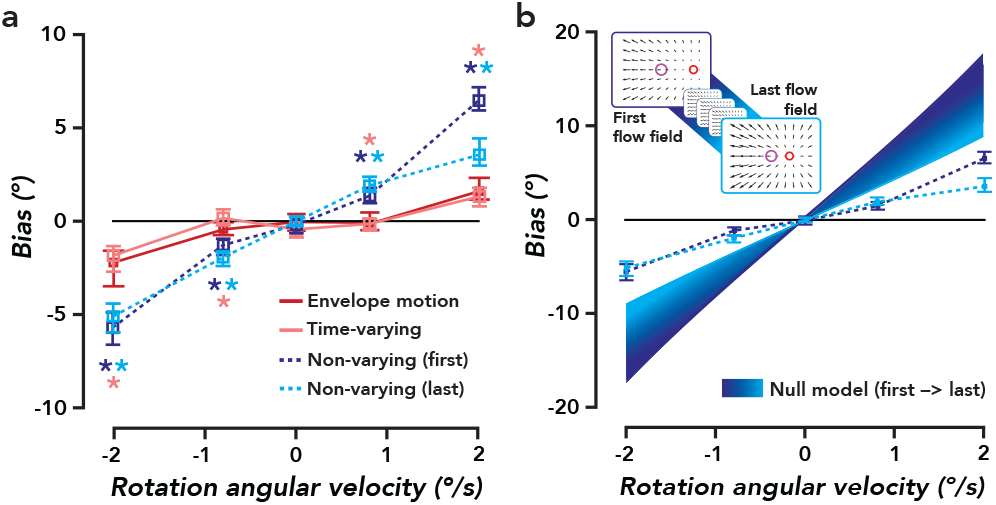
Bias in heading perception. **a.** Heading bias, pooled across observers (N observers = 10, N trials per rotation velocity per stimulus condition > 960). Square plot symbols, bias estimates. Error bars, 95% credible intervals. Asterisks represent statistical significance at a corrected cutoff of **α** = 0.04 (one-tailed permutation tests). The color of the asterisks indicates the specific hypothesis test that was performed. Light-blue asterisks, non-varying phase motion (last) was compared to envelope motion. Dark-blue asterisks, non-varying phase motion (first) was compared to envelope motion. Pink asterisk, non-varying (first) and non-varying (last) were compared to time-varying phase motion (an asterisk is plotted if both comparisons were significant). **b.** Heading bias for non-varying phase motion, pooled across observers (replotted from panel a). Color gradient, null model predictions across the sequence of optic flow fields and rotation velocities, indicating worst possible performance (upper bound on bias). The inset depicts the correspondence between the color gradient (from dark to light blue) and the sequence of optic flow fields.

Taken together, these results suggest that most observers relied on a heading perception algorithm that was more sophisticated than the null model, even when they only had access to instantaneous optic flow.

### Internal replication of non-varying phase motion rules out training duration and testing order effects

We conducted an internal replication of non-varying phase motion (first) after observers performed time-varying phase motion to validate that the observed reduction in bias was not simply due to days of exposure to stimuli utilizing phase motion. Indeed, bias for non-varying (first, initial) and non-varying (first, replication), were statistically indistinguishable for ±2 °/s and rightward 0.8 °/s rotations, when pooled across observers (*SI Appendix*, Fig. S3, Table S4). For leftward 0.8°/s rotation only, there was a small but significant difference between the two — an effect driven by O2 and O3 (Fig. S3). Together, we interpret this to mean that time-varying optic flow was necessary to achieve good heading perception, and this effect was not the result of training duration or the order of experimental conditions. We pooled non-varying (first) data — initial and replication — for all other analyses.

## Discussion

We developed a stimulus that conveys either an evolving sequence of optic flow fields (time-varying phase motion) or a single instantaneous optic flow field, the first or last in the sequence (non-varying phase motion). We found that heading judgments were strongly biased in the direction of rotation for non-varying, but nearly veridical for time-varying phase motion (i.e., less than 2° on average, below the accuracy required to avoid obstacles when moving at normal speeds (29, 30)). These effects were observed despite feedback indicating the correct heading direction following each trial (see *Methods*). That is, observers could not learn to make accurate or precise heading judgments without time-varying optic flow, even with extensive practice. This suggests that instantaneous optic flow is insufficient for heading perception in the presence of rotation and that time-varying optic flow is necessary. Stimulus duration was held constant across conditions, so these results cannot be attributed to differences in the duration of evidence accumulation. One implication is that nearly all models of heading estimation are incorrect, because they rely on instantaneous optic flow (4, 9, 19, 23, 24, 57, 58).

We propose a testable hypothesis, the ‘optic acceleration hypothesis’, for how observers use time-varying optic flow to generate rotation-tolerant estimates of heading (Fig. 1f, g), on the basis of a simple and general observation. Optic flow depends on observer translation and rotation, as well as depth structure (see *SI Appendix,* Eqs. 7, 18-20). When (retinocentric) heading and rotation velocity are constant over time (e.g., motion on a circular path), optic flow evolves solely because of changes in distance to objects in the environment (Fig. 1e, f; see *SI Appendix*). Under these conditions, heading is aligned with the singularity of the optic acceleration field (or “optic acceleration”; Fig. 1g; see *SI Appendix* for derivation). Thus, an estimator locating the singularity of optic acceleration would achieve unbiased performance. If heading and/or rotation velocity change over time, then such an estimator is biased, and the amount of bias depends on the magnitude of the change. The optic acceleration hypothesis is consistent with our experimental results. For time-varying phase motion, the optic acceleration field radiated from the heading direction at every moment in time. For non-varying phase motion, the optic acceleration field was by definition zero everywhere, hence it was completely uninformative. Heading bias was larger for non-varying than time-varying phase motion, for all observers.

According to the optic acceleration hypothesis, the precision of heading estimates should scale (inversely) with distance, or equivalently, with translational speed. The reliability and robustness of any algorithm for computing heading from optic acceleration depends primarily on d*p*/d*t*: the change in the inverse depth map over time (see *SI Appendix*). Previously published psychophysical results provide only indirect tests of this prediction, and the results are mixed. The proper manipulation to test our hypothesis — fixing rotation speed and varying translation speed, given constant rotation and heading — has not been performed. In one study (10), using a similar stimulus and task, the rotation and translation rates were simultaneously increased such that their ratio (59) was held constant. They found a non-significant trend such that the reliability of heading judgments increased with translation speed. In another study (15), rotation speed was fixed while translation speed was varied. They did not find significant differences in reliability across translation rates. However, observers were instructed to adjust the line-of-sight during each trial to estimate heading. Doing so introduced changes in rotation and (retinocentric) heading direction over time, both of which should cause errors under our hypothesis. Furthermore, the depth structure in this study was a dot cloud, providing rich parallax cues. Future studies attempting to test our hypothesis should eliminate all cues for heading other than time-varying optic flow (i.e., using time-varying phase motion and a single fronto-parallel plane), fix rotation speed while varying translation speed and/or manipulate distance, all while simulating constant observer translation and rotation (i.e., a circular path).

Previously published results demonstrate that heading bias depends on distance and speed. For example, one study (43) reported larger heading bias, approaching the optic flow singularity, when translation speed was lower (smaller d*p*/ d*t*), for translation plus a simulated eye movement with respect to a single fronto-parallel plane. Likewise, heading bias was larger when the initial distance to the plane was larger (smaller d*p*/d*t*). The optic acceleration hypothesis makes a direct prediction about precision, but not bias. Even so, we speculate that observers may fall back on a different (biased) strategy when faced with unreliable acceleration information (i.e., when d*p*/d*t* is very low), in the absence of other cues.

There is technically only one degenerate case for the optic acceleration hypothesis: travel parallel to a ground plane, when the inverse depth map does not change at all over time. Unexpectedly, heading performance can be good in such a scenario (38, 60), however interpretation is complicated because ground plane stimuli can convey an additional “horizon cue” (35, 61) as well as parallax. Even so, dramatic impairments in heading perception for travel parallel to a ground plane have been reported (62). Furthermore, including vertical objects (“posts") on a ground plane improves both passive path judgments (32) and active steering (40). The optic acceleration hypothesis may explain these results because inverse depth to the posts (but not the ground plane) changes over time, contributing to the optic acceleration field.

Previous models and computer vision algorithms have considered the importance of depth structure (variation in depth throughout the image at each instant in time) for heading estimation (4, 9, 19, 23, 57). According to the optic acceleration hypothesis, depth variation *per se* is not necessary for accurate heading perception, but richness of depth structure may improve the precision of heading judgments. This prediction is supported by experimental results. Observers can estimate heading nearly veridically (~1° of bias) for approach perpendicular to a single fronto-parallel plane (FOV = 90° x 90°) with a simulated eye movement (43). In a physiology experiment using a similar single-plane stimulus, VIP neurons signalled heading in a rotation-tolerant manner (i.e., less biased than the optic flow singularity) (46). In both cases, depth variation (and hence motion parallax) was virtually absent, but time-varying optic flow was present.

Increasing field of view (FOV) can enhance heading perception (32, 41–43, 59). According to the optic acceleration hypothesis, increasing FOV will lead to larger acceleration vectors in the periphery, conferring greater reliability in localizing the singularity of optic acceleration, and more precise heading estimates. Increasing FOV may reduce the small heading biases that we (and others) have observed for time-varying stimuli (envelope and time-varying phase motion).

Stereoscopic depth should enhance the reliability of d*p*/d*t*, leading to less variable and/or less biased heading estimates, under our account. This is borne out in human behavior (37, 38, 63–65) and might be mediated by the responses of neurons that signal heading and exhibit tuning for both binocular disparity and global motion patterns (66–69). The small heading biases that we (and others) have observed may be reduced when stereo cues are also available.

Human behavior and neurons in visual cortex are sensitive to optic acceleration. Biphasic temporal responses to motion have been measured psychophysically (70) and in MT neurons (71), suggesting that the temporal derivative of image velocity is computed. Global motion sensitive neurons that pool over local motion signals may inherit this property. Additionally, neurons in MT are sensitive in aggregate to acceleration, despite the sensitivity of individual neurons being low (72, 73). Likewise, psychophysical experiments reveal that humans are sensitive to optic acceleration (74), particularly in global motion patterns (75), with a frequency response peaking at around 1 Hz (76). Our stimulus contained primarily slow accelerations, well within the limits of sensitivity. One seminal study claimed that acceleration is not used, because they found comparable performance for 2-versus 3-frame dot lifetimes (31). However, the stimuli in that study simulated a circular path of travel parallel to a ground plane, the one case in which the (inverse) depth map does not evolve, and hence there was no optic acceleration in any of their stimuli.

We used non-varying phase motion to convey a single optic flow field. An ostensibly simpler approach would be to present only two frames of a dot motion stimulus. However, in practice, this would profoundly limit evidence accumulation and introduce artifacts (spreading the spatiotemporal energy) in the stimulus, thereby hindering performance. An alternative approach that has been employed is to present a video with many frames, but with each dot being presented for only 2 frames and then re-spawned at a random new location (31). Other studies have used dot lifetimes of hundreds of milliseconds (15, 33, 38, 42, 43, 55, 56). These methods remove or reduce image point trajectories, as well as accelerative image motion within a dot lifetime. They do not, however, remove time evolution of optic flow occurring *across* dot lifetimes (31). Although the observer does not have access to the continuous trajectory of each individual dot throughout the trial, he/she does have access to the continuous evolution of optic flow (see Fig 1f); its global structure evolves lawfully over time (7, 46, 52, 77) (see *SI Appendix*). Indeed, both human behavior (15, 31, 33, 78) and MSTd neurons that signal heading (55, 56) are sensitive to the global structure of the optic flow field, irrespective of particular dot positions. In summary, simply limiting dot lifetime does not remove time-varying optic flow, but non-varying phase motion does.

Time-varying phase motion eliminates looming, streamlines, dynamic perspective distortions and any other cue conveyed by image point trajectories and their spatial compositions (52, 53). Hence, our results suggest that these are not essential cues for heading perception. It has been proposed that dynamic perspective distortions caused by rotation (e.g., increasing retinal distance between points composing an object edge) help achieve rotation-tolerant heading estimates in macaque VIP neurons (46, 54) in the absence of motion parallax. Our finding of little bias for time-varying phase motion contradicts this conclusion and instead suggests that heading-tuned VIP neurons may be sensitive to optic acceleration.

Even so, many sensory cues may contribute to heading perception: motion parallax (10, 32, 42, 79), depth cues (e.g., from accommodation/blur, shading, looming, stereopsis) and extra-retinal signals (e.g., eye and neck efference copy (5, 8, 34–36, 44–47, 49, 80), vestibular (81, 82) and proprioceptive inputs (36)) could all play a role. The visual system seems to exploit available information, weighting signals according to their reliability, as has been shown extensively in the cue combination literature (83, 84). The small biases that we (and others) observed for time-varying stimuli may be abolished when these various cues are all consistent. Indeed, the simulated curvilinear motion stimuli in the current study may be viewed as a conflict between visual and vestibular signals. The resulting biases might, therefore, reflect an optimal combination of these two sources of heading estimates.

We have shown that time-varying optic flow is *necessary* for accurate heading perception in the presence of rotation, but additional experiments will be needed to test if it is *sufficient*. In our study, parallax (i.e., between near and far planes) was present in all stimuli. Measurements of bias for time-varying phase motion with an environment consisting of a single fronto-parallel plane (i.e., no local motion parallax) would reveal whether time-varying optic flow is sufficient. This seems plausible given that heading judgments can be nearly invariant to rotation for a single fronto-parallel plane with a large FOV (43), and that image point trajectories do not appear to be necessary for heading perception (15, 31, 33).

## Methods

### Participants

Data were acquired from eleven observers (seven males, four females). All observers were healthy adults, with no history of neurological disorders and with normal or corrected-to-normal vision. All but one (O8: the author C.S.B.) were naïve to the purposes of the experiment. Two observers, O1 and O9, had no prior experience participating in psychophysical experiments. The remaining observers had considerable experience with psychophysics, but little to none with heading perception tasks. One observer (O11) did not understand the task and was removed from the experiment following training. Observers O1–5 performed non-varying phase motion (first, initial), non-varying phase motion (first, replication), and time-varying phase motion. Observers O2 and O6–8 performed envelope motion. Observers O2 and O6-10 performed non-varying phase motion (last). Experiments were conducted with the written consent of each participant. The experimental protocol was approved by the University Committee on Activities Involving Human Subjects at New York University.

### Stimuli

There were three classes of stimuli: envelope motion, non-varying phase motion, and time-varying phase motion. Each of these stimuli comprised a field of plaid patches (Fig. 2). Each patch was the sum of two orthogonal gratings (spatial frequency, 3 cycles/°; contrast, 100%) multiplied by an envelope. The envelope was circular (diameter, 0.3°) with raised-cosine edges (width, 0.15°). No patches were presented within ±2.25° of the horizontal meridian unless they were beyond ±15° of the vertical meridian ensuring that observers could not directly track the singularity of optic flow, and so that fixation was not disturbed by the motion of the patches. The initial location of each patch was drawn from a uniform distribution. The initial phase of each grating was drawn from a uniform distribution at each trial’s onset. In all conditions, the stimulus was a movie with a duration of 4 seconds, presented with a refresh rate of 75 Hz on a calibrated HP p1230 CRT monitor (HP 2006) with a screen resolution of 1152 x 870 pixels, subtending 40° by 30°. Observers sat in a darkened room and viewed stimuli binocularly (but not stereoscopically) from a distance of 0.57 m, with their head in a chin/forehead rest to minimize head movements. Observers fixated a red dot (diameter, 0.09°) positioned centrally on a gray background throughout each stimulus presentation. Stimuli were generated and presented using MGL, a MATLAB toolbox for running psychophysics experiments (85).

For envelope motion, each patch (both the envelope and the plaid) was displaced on each frame (see *Movies S7-8*). The location of each patch corresponded to the projection of a point in a simulated 3-D environment onto the 2-D image plane (Fig. 1a). At each moment in time, the simulated cyclopean viewpoint of the observer (hereafter referred to as the “viewpoint”) moved according to the observer’s instantaneous translation and rotation. The locations of points in the 3-D environment and their projected 2-D locations were recomputed, and the patches were re-rendered at new locations (Fig. 1e). The phase of each plaid was randomly initialized and fixed with respect to its envelope on each trial. Envelope motion is equivalent to “dot motion” — a typical self-motion stimulus in the perceptual psychology literature (5, 10, 15, 31). As was the case with all of our stimuli, the retinal size of the plaid patches was constant over time, eliminating looming of individual elements as a cue.

The simulated 3-D environment comprised two rigid planes, initially 12.5 m and 25 m, respectively, from the viewpoint at the beginning of each trial (Fig. 1e). Each plane was composed of plaid patches. At the beginning of each trial, the density of patches in each plane was 0.16 patches/deg^2^, giving a total of ~168 visible patches forming each plane. Occlusion between planes occurred over time, but only for envelope motion. An additional 15° of padding with the same density was included beyond the field of view on each side to prevent any edges from appearing over time as the planes moved with respect to the viewpoint. The locations of plaid patches in the simulated 3-D space were projected onto the 2-D image plane using perspective projection (focal length, 0.57 m, i.e., equal to the viewing distance).

The simulated observer translated within a plane formed by the x and z axes, while rotating about the y-axis and looking down the z-axis (Fig. 1a). The path of movement was a circle and the line-of-sight (i.e., direction of gaze) was perpendicular to the planes at the beginning of each trial (10). The parameters of the simulated movement were ecologically valid. The chosen translation speed of 1.5 m/s corresponded to average human walking pace and the rotation speeds corresponded to movement along circles of radius 43 m (2 °/s rotation) or 107.4 m (0.8 °/s) or a straight path (0 °/s, circle of infinite radius). These circles are similar in radius to those used in previous studies (10, 15). The angle between the line-of-sight and heading direction was fixed within each trial, and was varied adaptively across trials. The line-of-sight corresponded to the center of the CRT monitor, where the fixation dot was located, and observers were told this. Eye tracking was used in all experimental conditions to monitor fixation (*SI Appendix*, *Eye Tracking*, for details).

For non-varying and time-varying phase motion, the plaid envelopes remained stationary while the phases of the gratings shifted over time. Each plaid comprised two orthogonal gratings, with component phase speeds specifying a 2-D velocity (86), giving a sampling of the underlying optic flow field at a fixed set of image locations. The phase velocity of the field of plaids corresponded to either a single optic flow field (“non-varying”) or a sequence of optic flow fields (“time-varying”). These optic flow fields corresponded to the instantaneous motion of points in the simulated 3-D environment relative to the observer. In the non-varying condition, the phases of a plaid shifted continuously throughout the trial according to the corresponding velocity vector within a single optic flow field (34). The phase change between successive frames was constant, so that the local velocity of each patch (and the optic flow field as a whole) was constant over time. The phase change of the vertical grating determined the horizontal component of the phase motion velocity, while the phase change of the horizontal grating determined the vertical component of the velocity. For each (vertical or horizontal) grating, the phase change between frames was computed from the corresponding (horizontal or vertical) component of the optic flow velocity: 2π times the product of the (horizontal or vertical) speed (°/s) and the spatial frequency (cycles/°), divided by the elapsed time (1/s).

On each trial of non-varying phase motion, a video was presented that corresponded to a particular instantaneous optic flow field (see *Movies S1-4*). The optic flow field presented was either the first or last in the same time-varying sequence used in the envelope motion and time-varying phase-motion conditions (Fig. 4b inset). We called these stimuli “non-varying phase motion (first)” and “non-varying phase motion (last),” or more simply: “non-varying (first)” and “non-varying (last).”

In the time-varying phase motion condition, the phase change between successive frames varied throughout the trial, conveying a sequence of optic flow fields (see *Movies S5-6*; *Fig. 1f*). The phase change between frames was computed by generating a sequence of 10 optic flow fields spaced evenly in time over the 4 second trial and calculating the phase change for each optic flow field. These phase changes, which we defined as “time-varying optic flow,” were determined by the movement of the viewpoint relative to the environment, just as in the envelope motion condition (see *SI Appendix* for details). Time-varying optic flow consisted of local accelerations and decelerations of the phase motion. Consequently, time-varying phase motion conveyed a sequence of optic flow fields temporally subsampled from the full sequence conveyed with envelope motion. The crucial difference, however, between these stimuli was that the patch envelopes did not move at all in the former, but did move in the latter. The trajectories of individual image points may provide a cue for computing heading. Phase motion (time-varying or non-varying) eliminates this cue. If you walked on circular path with a perforated cardboard board (with many small holes) affixed to your head a few inches out, you would see the visual image evolve only through those apertures. If optic flow estimation were possible in each aperture, this scenario would be analogous to time-varying phase motion.

To generate phase motion stimuli, we computed the optic flow field(s) corresponding to the motion of the observer’s viewpoint relative to the simulated 3-D environment for specific combinations of heading and rotation velocity (*SI Appendix*, Eqs. S1–9). Optic flow for a planar depth structure can be expressed in closed form. Thus, we simply evaluated this function at the x and y image locations of the plaid patches, for each plane separately. A more general approach that will work for analytically intractable depth maps is to densely sample the depth map, compute an equally dense optic flow field from it, and finally subsample the optic flow field at specific screen locations. These calculations were implemented in custom MATLAB software.

### Protocol

Observers viewed videos of plaid patches whose image motion corresponded to movement along a circular path with variable combinations of heading direction and rotation velocity, and subsequently performed a forced-choice heading discrimination. This task, with the envelope motion stimuli, nearly replicated a previous study (10), and we refer the reader to their methods and figures for elaboration. The main differences between our study and theirs were: 1) we provided correct/incorrect feedback after each trial and they did not; 2) their translation rate was 2 m/s and ours was 1.5 m/s; 3) our stimuli consisted of circular plaid patches and theirs consisted of dots; and 4) our stimulus movie was 4 seconds in duration and theirs was 2.33 seconds. The average heading biases measured in that study were within one degree of ours, suggesting that, following training, biases are present with or without tone feedback (for time-varying stimuli), also consistent with the results of a perceptual learning study (48). Movement along a circular path avoids ambiguity about which coordinate system the heading judgment was made in (because retinocentric heading is constant over time) (10). Using a retinocentric judgment helps insure against the possibility of heading estimates being influenced by path perception (10, 15, 33).

Each trial consisted of a 4 second video presentation followed by a 1 second inter-trial interval (ITI), during which the observer made a key press response indicating whether they perceived their heading to be left or right of center (i.e., the fixation point). Observers rarely failed to respond during the 1 second ITI (5 out of 30,000 trials). Immediately following key press response, observers were given auditory feedback indicating if their response was correct or incorrect (two different tones). The ground truth heading on each trial was set adaptively using two interleaved one-up-one-down staircases (per rotation speed) that converged to 50:50 leftward:rightward choices (i.e. the heading equally often judged as left and right of center). Staircases (initial step size, 4°) were initialized with a heading of ±20° every 30 trials (per staircase) to collect sufficient data at the asymptotes and center of the psychometric function (Fig. 3). There were 6 staircases in total running simulataneously, and they continued across blocks of trials.

The order of sessions (test and training inclusive) was: envelope motion, non-varying (first, initial), time-varying, non-varying (first, replication), non-varying (last). One participant (O2) performed this whole sequence. One subgroup (O6–8) performed only envelope motion and non-varying (last). Another (O1, O3–5), non-varying (first, initial), time-varying, and non-varying (first, replication). Another (O9-10), just non-varying (last). These subgroups were tested in the same order, given that constraint. Each experimental condition consisted of two test sessions (plus prior training sessions), each session comprised 10 blocks, and each block comprised 60 trials, for a total of 600 trials per session. Each session was around an hour in length, including breaks, and was conducted on a different day. Observers were allowed to take breaks between blocks of trials to prevent fatigue. In all conditions, the observer’s translation speed was fixed at 1.5 m/s, and the angular velocities of the simulated rotations were −2 °/s, −0.8 °/s, 0 °/s (“no-rotation”), +0.8 °/s, or +2 °/s. Negative signed velocities are leftward rotation and positive velocities are rightward rotation. These five rotation velocities were interleaved in a randomly permuted order, ensuring equal numbers of trials with each velocity within each block of trials.

Prior to encountering each new stimulus condition, observers performed a minimum of 120 training trials (identical to experimental trials). Training always began with minimum of 60 trials without rotation, followed by a minimum of 60 trials with rotation. Training ensured that observers understood the protocol and had reached asymptotic perceptual sensitivity in the task. One observer required only 120 trials of training in each condition (O5), but most required one or two hour-long training sessions (600 trials each) to achieve asymptotic performance on the task, consistent with previous reports (10, 48). To determine when to end training and start the experiment, we checked that that discrimination accuracy was ~70-80% across all trials and ensured that the average converged heading values were consistent across the two interleaved staircases (for each rotation speed) and stable across runs of trials. We also visually inspected the staircases to see that they had converged properly. Observers’ discrimination thresholds decreased over training, asymptoting at 1° to 8°, depending on the stimulus and rotation velocity (*SI Appendix*, Fig. S2). Observers were never given feedback about rotation speed or information about the number of rotation speeds tested during training or the experiment.

### Data Analysis

Heading bias and discrimination threshold were estimated using Psignifit 4, a MATLAB toolbox for Bayesian inference for psychometric functions (87). Cumulative normal psychometric functions were fit separately to each individual observer’s responses and also to the pooled responses of all observers. We fit a beta-binomial model with four free parameters: *μ*, σ, λ, η, which respectively corresponded to the mean and slope of the psychometric function, the lapse rate, and the over-dispersion of the data relative to the binomial model (Fig. 3). The mean of the psychometric function or PSE describes the heading for which the observer reported 50:50 leftward:rightward of center (i.e., leftward:rightward of the location of the fixation point). A *bias* of 0° minus this heading is needed to cancel out the horizontal shift in the psychometric function. Bias describes how far the observer’s internal estimate of center is from 0° (i.e., true center). For example, if an observer viewed a pure translational (0° heading) optic flow field and had a bias of −2°, that would suggest that the observer would interpret a heading of 0° as a leftward heading of 2°. The slope of a cumulative normal psychometric function determines the discrimination threshold of the observer in a 2-AFC task, defining how sensitive of a classifier they are (*SI Appendix*, Fig. S2). Lapse rate describes how often observers lapsed, pressing the wrong key by accident. It sets an upper limit on accuracy by jointly controlling the psychometric function’s upper and lower asymptotes. Over-dispersion describes how much the dispersion of the observed data exceeded the dispersion expected under a binomial model, potentially due to serial dependencies, changes in arousal throughout the experiment, or other factors. The likelihood of the data given the model was evaluated, and flat priors were used for each of the four free parameters, yielding a posterior describing the probability of the model given the data. We report maximum a posteriori (MAP, equivalent in this case to maximum likelihood) estimates for each parameter by numerical integration over the 4-D posterior, p(**ϴ**|data), where **ϴ** is a vector containing the four free parameters, and finding the maximum of the marginal distribution for each parameter. 95% credible intervals for parameter fits were computed as the central 95% density region of the cumulative probability function of each marginal posterior distribution. Statistical comparison of heading biases across conditions was performed via permutation test (see *SI Appendix*, *Statistics,* for details).

### Null Model

When there is observer rotation in addition to translation, the singularity of the optic flow field is displaced away from the heading in the direction of rotation (Fig. 1b–d). Thus, an estimator using only the optic flow singularity would be systematically biased — reporting heading as the visual angle subtending the singularity location. We defined our null model in this way (as in Stone and Perrone, 1997 (10)). The singularity position was computed by extracting the horizontal component of optic flow (see *SI Appendix*, Eq. S7) and solving for the point at which this function crossed zero. Null model predictions were generated for the near (12.5 m) plane across a finely spaced range of rotation angular velocities (−2 to +2 °/s) and times (0–4 seconds of the trial), using the same parameters that were used to generate the experimental stimuli. Null model predictions for the far plane were much larger than for the near plane (i.e., around twice as large) and were not reported because observers always performed much better than that. The null model is not a serious model of how the visual system computes heading under natural conditions. It is a biased algorithm by construction — a lower bound on performance we are using as a reference point for the observed heading biases in the non-varying phase motion condition.

## Acknowledgements

Special thanks to Shani Offen, Davis Glasser, Elisha Merriam, and Ionatan Kuperwajs for conducting preliminary versions of these experiments. Thanks to Shannon Locke, Hörmet Yiltiz, and Emmanouil Protonotarios for providing comments on experimental design and analysis. Thanks to Greg DeAngelis and Eero Simoncelli for helpful discussion. Thanks to Michael Landy, Bas Rokers, and Kathryn Bonnen for providing invaluable feedback on the manuscript. This research was supported by the National Eye Institute Visual Neuroscience Training Grant, T32 EY007136 (to C.S.B. through NYU) and the National Defense Science & Engineering Graduate Fellowship (to C.S.B.).

## Author Contributions

Conceptualization D.J.H. and C.S.B.; Investigation, C.S.B.; Analysis, C.S.B.; Software and visualization, C.S.B.; Writing, C.S.B. and D.J.H; Supervision, D.J.H.

## Supplementary Information

### Supplementary Methods

#### Derivation of the optic flow field in terms of observer translation and rotation

Points on the surface of objects in the virtual 3-D environment are represented as position vectors **X** = *(X, Y, Z)*^T^ relative to a viewer-centered (i.e., retinocentric) coordinate frame. When the viewpoint moves, this point moves along a 3-D path **X**(t) = *(X(t), Y(t), Z(t))*^T^ and the instantaneous velocity of this point in 3-D is the derivative of this path with respect to time:

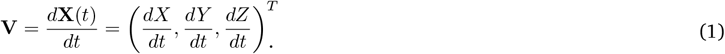

The 3-D points project to image locations. As each 3-D point moves over time, its corresponding image location changes over time. Under perspective projection, the point **X** projects to the image point *(x, y)*^T^ given by:

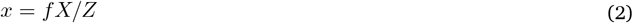

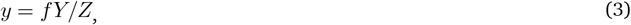

where f is the focal length of the projection. As the 3-D point moves over time, its corresponding 2-D image point traces out a 2-D path *(x(t), y(t))*^T^, the gradient of which is the image velocity:

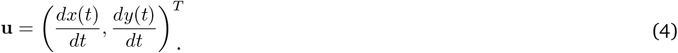

Combining Eqs. 2, 3, and 4 gives an expression for image velocity in terms of the 3-D position and velocity of the object surface point:

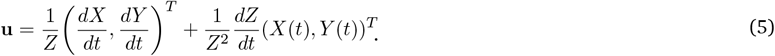

If one considers the paths of all visible 3-D surface points, and their projections onto the image plane, then one obtains a dense set of 2-D paths, the temporal derivative of which is a vector field of 2-D velocities, commonly known as the optic flow field.

We assume that the objects in the scene move rigidly with respect to the viewpoint, as though the simulated observer were moving through a stationary environment. This is a special case because all points on a rigid body share the same six motion parameters relative to the viewer-centered coordinate frame. In particular, the instantaneous velocity of the observer through a stationary scene can be expressed in terms of the observer’s 3-D translation **T** = (*T*_x_, *T*_y_, *T*_z_)^*T*^, and his/her instantaneous 3-D rotation **Ω** = (*Ω_x_*, *Ω_y_*, *Ω_z_*)^*T*^. Here, the orientation of **Ω** gives the axis of rotation, while |**Ω**| is the magnitude of the angular velocity. Given this motion of the viewpoint, the instantaneous 3-D velocity of a surface point in viewer-centered coordinates is:

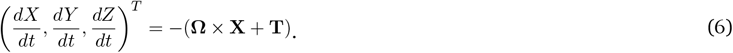

The 3-D velocities of all surface points in a stationary scene depend on the same rigid-body motion parameters, and are given by Eq. 6. It has previously been shown that if one substitutes these 3-D velocities for d**X**/d*t* in Eq. 5, an expression for the form of the optic flow field for a rigid scene can be obtained (1):

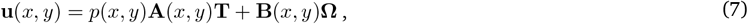

where *p(x, y)* = 1/*Z(x,y)* is inverse depth at each image location, and

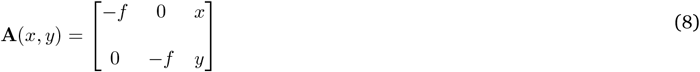

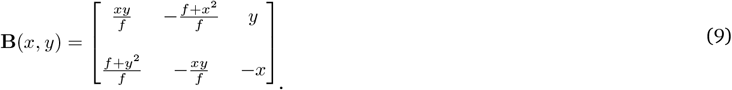

The matrices **A**(*x,y*) and **B**(*x*,*y*) depend only on image position and focal length.

Equation 7 describes the flow field as a function of 3-D motion and depth. It has two terms. The first term is referred to as the translational component of the optic flow field because it depends on 3-D translation and 3-D depth. The second term is referred to as the rotational component because it depends only on 3-D rotation. Since *p(x,y)* (the inverse depth) and **T** (the translation) are multiplied together in Eq. 7, a larger distance (smaller *p*) or a slower 3-D translation (smaller |**T**|) both yield slower image velocity. In a pure translation optic flow field, the direction and magnitude of each vector depends on the inverse depth at the 3-D location that projects to that vector’s corresponding location on the image plane (Fig. 1b). When the observer performs a pure translational motion, features in the image move toward or away from a single point in the image, called the focus of expansion (FOE). By contrast, the optic flow field’s rotational component (**Ω,** Eq. 7) does not depend on inverse depth or scene structure. Therefore, rotation in the absence of translation results in a quadratic flow field, that is, each velocity vector (*u*_x_, *u*_y_) is a quadratic function of image position (Eq. 7, Fig. 1c). In a translation + rotation optic flow field, the translational and rotational components are summed, and the optic flow singularity (point where image velocity is zero) no longer corresponds to the direction of translation (Fig. 1d).

The derivation above characterizes the optic flow field with respect to 3-D translation and rotation in a viewer-centered coordinate frame. The horizontal component of heading, *h_x_*, is defined as tan^−1^(*T_x_* / *T_z_*), the angle subtended by the translation vector (*T*_x_, *T*_z_)^*T*^ and the z-axis, in the viewer-center coordinate frame. For the case simulated in our experiment, movement along a circular path with the globes fixed in the orbits and the head fixed on the body, heading does not change over time. In this case, the viewpoint’s translation velocity **T** and rotation velocity, **Ω,** are constant over time, and there is a time-invariant angle between the translation vector **T** and “line-of-sight” (i.e., the orientation of the viewer-centered coordinate system). In our heading discrimination task, an ideal classifier comparing *h_x_* to zero (<0: “left”; >0: “right”) would achieve perfect performance.

#### Computing heading from the optic acceleration field

If **T**(*t*), and **Ω**(*t*) are (approximately) constant over time, then only the first term in Eq. 7 remains time-dependent. In that case, the optic acceleration field — the temporal derivative of Eq. 7 — is:

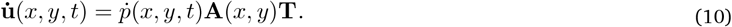

Overdots denote temporal derivatives. The optic acceleration field in Eq. 10 is similar to the translational component of the optic flow field — the first term in Eq. 7 — but is weighted by the temporal derivative of the inverse depth, d*p*/d*t*, instead of by *p*.

Next we solve for the singularity of optic acceleration — the image coordinates at which optic acceleration is zero — denoted *x*_0_ and *y*_0_. Given the definition of **A** in Eq. 8:

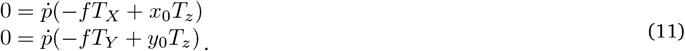

If the time derivative of the inverse depth *p* is non-zero, then

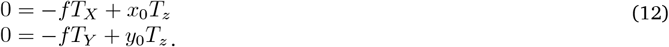

Thus the optic acceleration singularity is only a function of the instantaneous translation **T** and the focal length *f*:

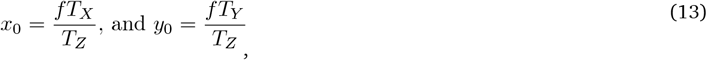

and the heading **h** is equal to the angles subtended by the image position of the optic acceleration singularity:

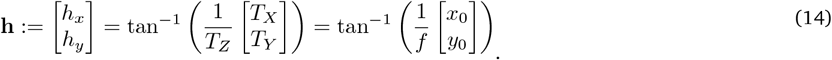

The assumption of constant **Ω**(*t*) and **T**(*t*) is consistent with the case simulated in our experiment — a circular path with a time-invariant angle between the translation **T** and line-of-sight — and is approximately true whenever observer motion changes slowly.

The reliability of heading estimation with this approach depends on non-zero changes in inverse depth over time. If d*p*/d*t* is zero (e.g., consistent with travel parallel to a ground plane), then Eq. 10 is degenerate and there is no way to solve for heading direction from Eq. 10. As d*p*/d*t* becomes smaller, the heading estimates will become unreliable in the presence of noise.

If, instead, **T**(t) rotates over time by a constant **Ω** (e.g., corresponding to making a constant velocity eye movement while travelling on a straight path), the optic flow field is:

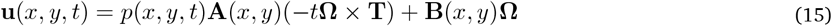

where **T** is the initial constant translation. For t>0, the optic acceleration field is:

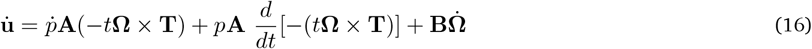

Solving for singularity of optic acceleration,

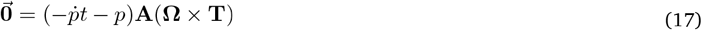

When *p = t dp/dt* and *Ω*_*x*_ *and Ω*_*z*_ = 0, we arrive again at Eq 13 and 14. Heading **h** is equal to the image position of the singularity of optic acceleration. Once heading is known, there are algorithms for estimating rotation and depth (1). Some of these observations about the mathematical relationships between heading, optic flow, and optic acceleration have been reported previously (2–7).

#### Components of time-varying optic flow

Travel parallel to a ground plane is the sole situation in which optic flow does not evolve over time. Although this case never occurs outside the laboratory, it has been simulated in many psychophysical experiments, and intuitions have been formed based on it. Here, we emphasize that in *all other scenarios*, the optic flow field evolves over time, in a manner depending on observer translation, rotation, and the depth structure of the environment. We will derive two simple cases to illustrate our point, but do not intend this to be a comprehensive treatment of this topic.

For pure translation at a constant heading (in a non-ground-plane environment), the optic flow field evolves over time because the angular subtense of surfaces in the environment changes (nonlinearly) with distance. This causes a change in the magnitudes of the velocity vectors, but preserves the angles. Thus, the optic flow singularity stays fixed over time (10). This is clear via the same logic as Eqs. 10–13. If **T**(*t*) is constant, **Ω**(*t*) is zero, it follows from Eq. 7 that:

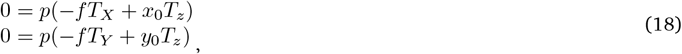

where *x*_0_ and *y_0_* are the image points corresponding to the singularity of the optic flow field. If *p*(*x*,*y*,*t*) is non-zero, we arrive at Eqs. 13–14: the singularity gives the heading, which does not change over time.

If the observer instead moves along a circular path with a fixed angle between the line-of-sight and the heading, that implies that **T**(*t*) and **Ω**(*t*) are non-zero and constant. In this case, the optic flow singularity moves over time, depending only on the evolution of the inverse depth map *p*(*x*,*y*,*t*), because the second term in Eq. 7 is a constant:

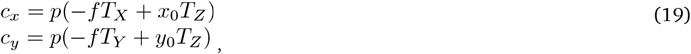

where **c** = (*c_x_*, *c_y_*)^T^ = **BΩ** is the constant rotation component (the second term) in Eq. 7. Thus,

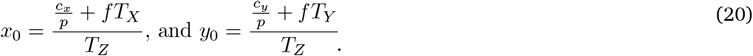

Note that the only term in the optic flow singularity that depends on time is *p*. For a simple depth structure like a plane or other geometric primitive, it is easy to see that the optic flow singularity will move smoothly over time.

If one moves the eyes and/or head independently of the body, **T**(*t*) will be non-constant. A simple example is movement along a straight path with a constant velocity rotation of the eyes or head (i.e., **Ω**(*t*) is constant). Here, **T**(*t*) changes over time only according to **Ω**. That is, heading drifts on the retina according to the rotation. In this case, the magnitude and angles of the optic flow vectors change over time because of coordinate system rotation, and also because the translational (but not rotational) component of optic flow (i.e., first term in Eq. 7) evolves due to the changing depth map.

To summarize, we have defined two components of time-varying optic flow: coordinate system rotation and change in the depth map over time due to movement relative to the environment. If the observer moves along a circular path with a fixed angle between the line-of-sight and the heading, only the second component remains, and the optic flow will evolve (and optic flow singularity will shift) over time depending only on the depth map. Thus, this special case, heading on a circular path, avoids ambiguity about which coordinate system the observer made their judgment in (because it is fixed over time). This was the case chosen for our psychophysical experiments.

#### Statistics

The strength of statistical evidence for differences in heading bias across conditions was assessed via permutation test, a resampling technique that constructs a null distribution of a test statistic from the observed data (8–9). The null distribution was constructed iteratively in the following way. We pooled the trial-to-trial responses across the two conditions of interest, randomly shuffled the condition labels (permutation without replacement), split the data into two groups (group size was determined by the number of trials collected in the corresponding condition), found the MAP estimate of *μ* (heading bias, see *Data Analysis* in the main body of the text) for each group, and computed the difference between the two groups’ heading bias estimates. This procedure was iterated 1000 times. The test statistic was the difference in heading bias across conditions. For the translation + rotation conditions (−2, −0.8, +0.8, & +2 °/s rotation), exact p-values for each comparison were calculated by finding the proportion of test statistics in the null distribution that were greater than (for rightward rotation) or less than (for leftward rotation) the observed difference in heading bias computed from the observed, un-shuffled data (one-tailed permutation test). For the no-rotation condition (0 °/s), the absolute values of the null distribution and observed statistic were used instead to perform a two-tailed permutation test. This procedure was carried out for every comparison except for that between envelope motion and time-varying phase motion, for which a two-tailed test was performed for each rotation velocity. This was because we did not hypothesize that one condition would have larger biases (see *Introduction* and *Discussion*). For the one-tailed tests, the direction of the test was determined by our predictions. See Tables S1-5 for the full set of predictions and corresponding exact p-values.

P-values were corrected for the finite number of permutations. The purpose of a permutation test is to estimate the null distribution of a test statistic, ideally enumerating each possible permutation and obtaining an estimate of the p-value, denoted *p*∞, under the null hypothesis. However, exhaustive permutation is often computationally intractable and instead, often only a random subset of possible permutations are performed, as was the case with this analysis. It has been shown that simply treating the computation of *p*∞ as an estimation problem by replacing *p*∞ with an unbiased estimator can lead to an inflated type I error rate.

Furthermore, p-values of zero are commonly reported in the literature, but these probabilities make little sense inferentially. If all permutations were enumerated, one would obtain the observed statistic, and the p-value would be greater than zero. Furthermore, in the context of multiple comparisons, p-values of zero become more perilous, and may lead to faulty inference. To partially correct for these issues, all p-values were subjected to a simple and well-known calculation that effectively estimates exact p-values:

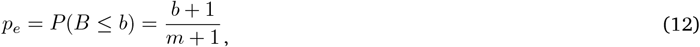

where *p_e_* is the exact p-value, *b* is the number of permuted test statistics greater than the observed test statistic, and *m* is number of permutations performed (10–11).

P-values were also corrected for multiple comparisons. 350 hypothesis tests were performed. We note that not all tests were independent, but that positive dependencies were expected to exist between them because they were computed using the same test statistics. To control the overall type I error rate at an alpha level of .05 while assuming dependencies between tests, we subjected all exact p-values to a “two-stage” adaptive procedure (12–13). It has been shown via simulation that of a number of similar tests, this adaptive test is best at controlling false discovery rate (FDR), regardless of effect size, in the presence of positive dependence (14). This procedure yielded a corrected critical p-value, **α** = 0.03996004 (rounded off to 0.04 in the main body of the text), which was adopted as a conservative cutoff to judge the strength of the statistical evidence for each hypothesis test.

#### Eye tracking

Measurements of gaze position were used to validate that observers were fixating the central red dot. Gaze position was measured using a remote infrared video-oculographic system (Eyelink 1000; SR Research), with a spatial resolution of 0.01° and average accuracy of 0.25-0.5° when using a head rest. Gaze position was acquired at a sampling rate of 500 Hz. Blinks were removed from the gaze-position time-series and the missing gaze positions were linearly interpolated. All observers fixated stably during the 4 second stimulus and mean eye position did not depend on the velocity of simulated rotation. The medians (across observers) of the average counts (across trials) of saccades per trial were: 1.1, 0.92, 0.82, 0.91, and 1.2 (SEM, 0.8705, 0.8762, 0.8302, 0.9263, and 0.8876) for −2, −0.8, 0, +0.8, +2 °/s rotations. Only observer O5 and O6’s saccade *frequency* (average across trials) was dependent on the simulated rotation velocity of the stimulus, but this dependence was small. For O5, the difference in the average number of saccades per trial between ±2 °/s rotations and no-rotation (0 °/s) was 0.4. For O6, this difference was 1. The remaining observers’ saccade counts showed little or no dependence on rotation velocity (difference in average number of saccades per trial between ±2 °/s rotations and no-rotation = 0.03, N = 10).

---

**Fig. S1.**
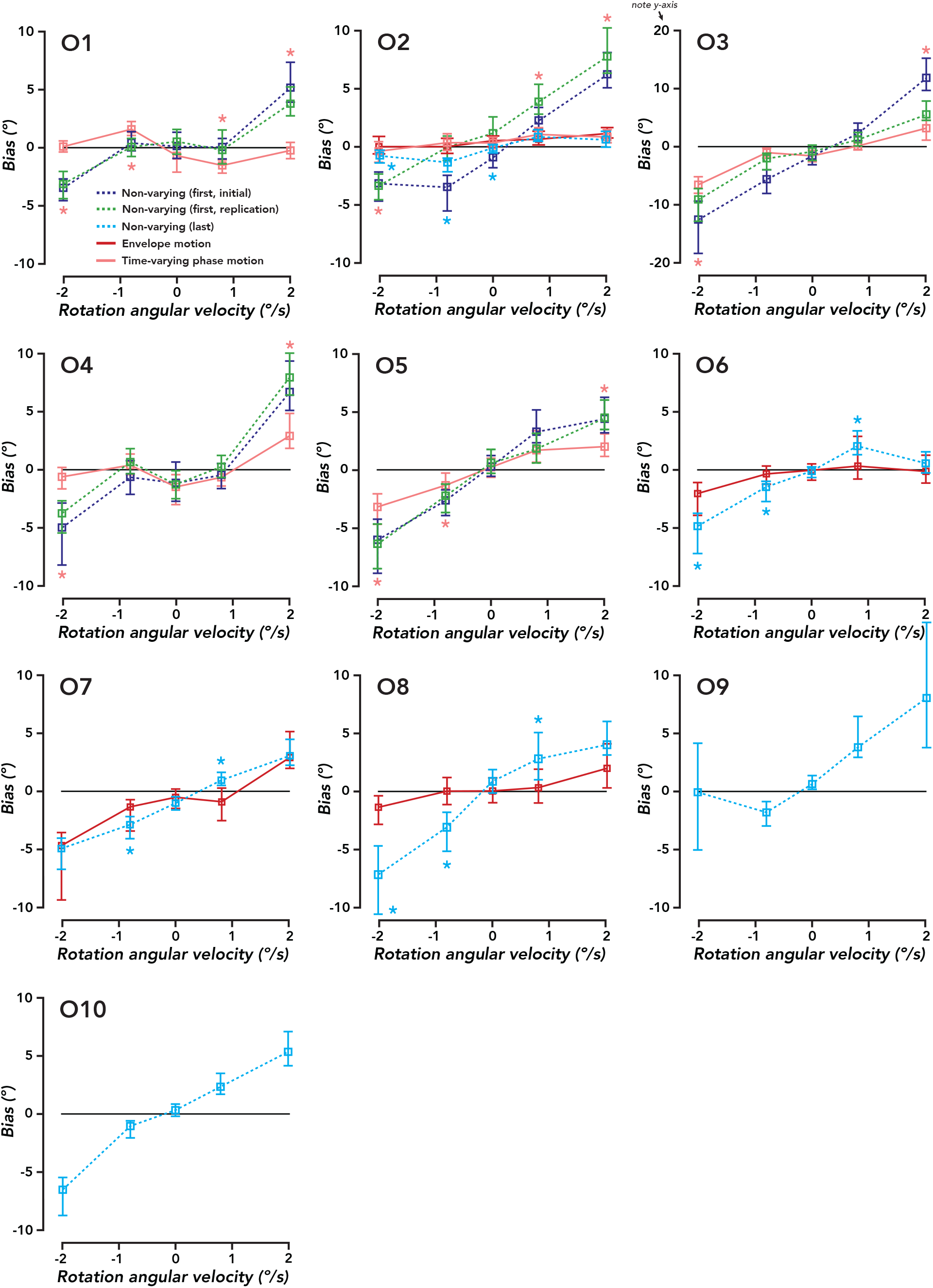
Heading bias for each observer. Same format at Fig. 4a. Heading bias for each individual observer. Square plot symbols, MAP estimates of bias. Error bars, 95% credible intervals. Asterisks represent statistical significance at a corrected cutoff of **α** = 0.04. The color of the asterisks indicates the specific hypothesis test that was performed. Pink asterisks, non-varying (first, initial) and non-varying (first, replication) were separately compared to time-varying phase motion (one-tailed permutation test). If both comparisons were significant, a pink asterisk was plotted. If only one comparison was significant, no asterisk was plotted (see Table S2 for all relevant p-values). Light-blue asterisks, non-varying (last) was compared to envelope motion (one-tailed permutation test).

**Fig. S2.**
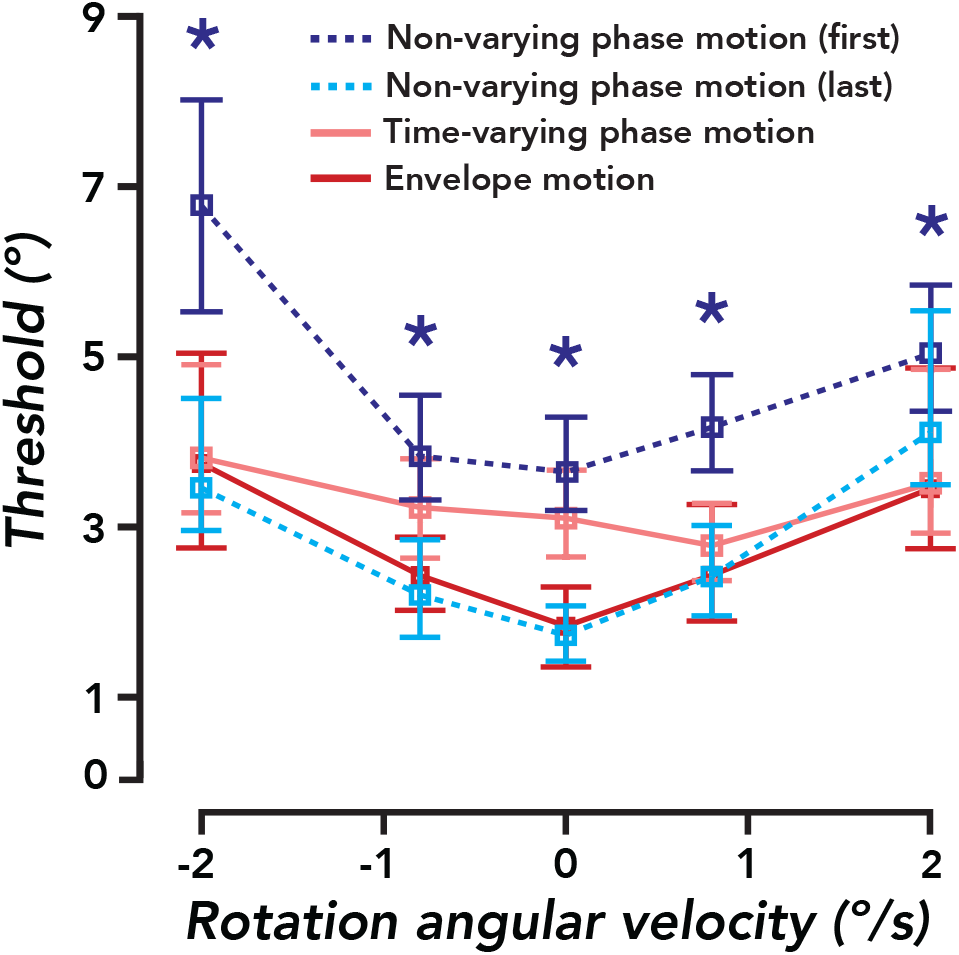
Heading discrimination thresholds. Analysis performed on the pooled data of all observers (N observers = 10). Each discrimination threshold was computed as the MAP estimate of σ, the slope parameter of a cumulative normal psychometric function. Square plot symbols, MAP estimates of σ. Error bars, 95% credible intervals. Asterisks represent statistical significance with a cutoff of **α** = 0.04. Dark-blue asterisk, non-varying (first, pooled over initial and replication) was compared to envelope motion (one-tailed permutation test). All depicted p-values, p < 0.013.

**Fig. S3.**
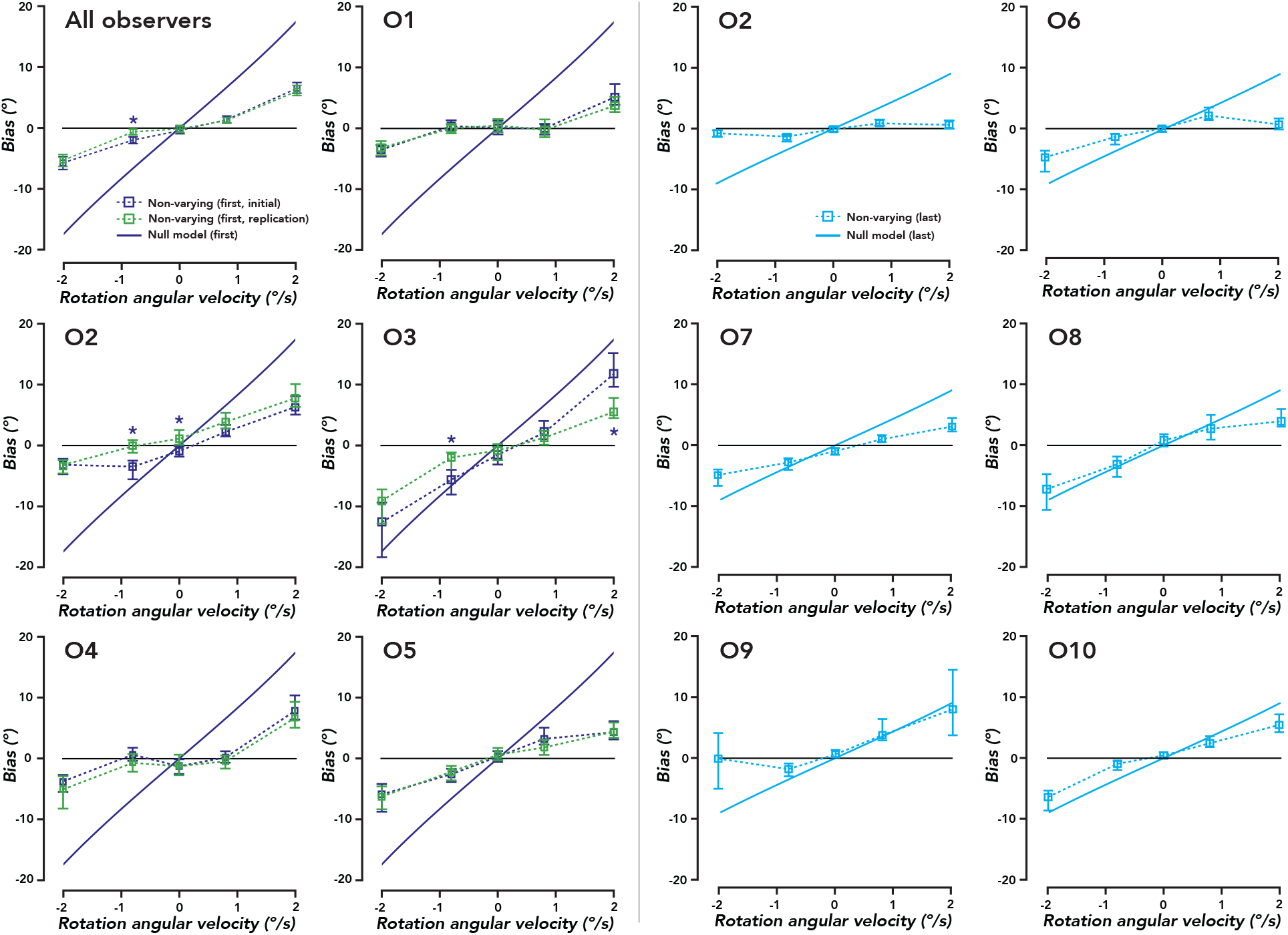
Heading bias for non-varying phase motion (for each observer) versus null model. Square plot symbols, bias estimates for non-varying phase motion. Solid lines, null model predictions. Dark-blue asterisks indicate that for the corresponding rotation velocity, absolute bias for non-varying (first, initial) was significantly larger than that for non-varying (first, replication). For most observers, heading bias fell between veridical performance and the prediction of the null model. The null model prediction fell within the 95% CI of the estimate of heading bias for some observers (O3, O6, O7, O8, O9), for a subset of rotation velocities. Note: non-varying (last) data (pooled, all observers) are plotted in Fig. 4b, so not replotted here.

**Table S1.** Strength of statistical evidence for hypothesis that instantaneous optic flow is insufficient for accurate heading perception in the presence of rotation. **a.** Non-varying (first, initial) > envelope motion. **b.** Non-varying (first, replication) > envelope motion. **c.** Non-varying (first, pooled data from **a** and **b**) > envelope motion. Corrected exact p-values for observer O2 (N trials per rotation velocity = 240) and pooled across observers (“All”; N observers = 10, N trials per rotation velocity > 960). **d.** Non-varying (last) > envelope motion (“All”; N observers = 6, N trials per rotation velocity > 960). One-tailed permutation tests were performed for −2, −0.8, +0.8, and +2 °/s rotation velocities, and a two-tailed permutation test was performed for the 0 °/s (no-rotation) condition. The tail of the test depended on the sign of the rotation velocity. For example, the hypothesis tests in panel **a** assessed whether the bias for non-varying (first, initial) trials was higher than the bias for envelope motion trials for +0.8 and +2 °/s rotations, and lower for −0.8 and −2 °/s rotations. This convention is used throughout all tables. All boldface p-values are less than than the critical exact p-value (cutoff) of **α** = 0.04 (see *Statistics* for details). P-values are rounded to three significant digits.

| **a** | -2 °/s | -0.8 | 0 | +0.8 | +2 | **b** | -2 °/s | -0.8 | 0 | +0.8 | +2 |
| --- | --- | --- | --- | --- | --- | --- | --- | --- | --- | --- | --- |
| All | **0.002** | **0.001** | 0.211 | **0.001** | **0.001** | All | **0.001** | 0.294 | 0.802 | **0.001** | **0.001** |
| O2 | **0.001** | **0.001** | **0.002** | **0.001** | **0.001** | O2 | **0.001** | 0.435 | 0.180 | **0.001** | **0.001** |
| **c** | -2 °/s | -0.8 | 0 | +0.8 | +2 | **d** | -2 °/s | -0.8 | 0 | +0.8 | +2 |
| All | **0.002** | **0.010** | 0.447 | **0.002** | **0.001** | All | **0.001** | **0.001** | 0.748 | **0.001** | **0.002** |
| O2 | **0.002** | **0.034** | 0.053 | **0.001** | **0.001** | O2 | **0.013** | **0.001** | **0.002** | 0.134 | 0.963 |
|  |  |  |  |  |  | O6 | **0.004** | **0.029** | 0.825 | **0.001** | 0.113 |
|  |  |  |  |  |  | O7 | 0.372 | **0.001** | 0.215 | **0.001** | 0.170 |
|  |  |  |  |  |  | O8 | **0.001** | **0.001** | 0.200 | **0.011** | 0.097 |

**Table S2.** Heading bias for non-varying phase motion > time-varying phase motion. Evidence that time-varying optic flow is necessary for heading perception in the presence of rotation. Same format as Table S1. Corrected exact p-values for each observer (N trials per rotation velocity = 240), and pooled across observers (“All”; N observers = 5; N trials per rotation velocity > 1200). **a.** Non-varying (first, initial) > time-varying phase motion. **b.** Non-varying (first, replication) > time-varying phase motion. **c.** Non-varying (first, pooled data from **a** and **b**) > time-varying phase motion. **d.** Non-varying (last) > time-varying phase motion.

| **a** | -2 °/s | -0.8 | 0 | +0.8 | +2 | **b** | -2 °/s | -0.8 | 0 | +0.8 | +2 |
| --- | --- | --- | --- | --- | --- | --- | --- | --- | --- | --- | --- |
| All | **0.001** | **0.001** | 0.965 | **0.001** | **0.001** | All | **0.001** | **0.005** | 0.323 | **0.001** | **0.001** |
| O1 | **0.001** | **0.002** | 0.045 | **0.019** | **0.001** | O1 | **0.001** | **0.009** | 0.211 | **0.001** | **0.001** |
| O2 | **0.007** | **0.002** | 0.421 | **0.001** | **0.001** | O2 | **0.001** | 0.204 | **0.016** | **0.001** | **0.001** |
| O3 | **0.008** | **0.001** | 0.285 | 0.066 | **0.001** | O3 | **0.035** | 0.107 | 0.274 | 0.049 | **0.023** |
| O4 | **0.015** | 0.050 | 0.660 | 0.339 | **0.003** | O4 | **0.004** | 0.672 | 0.753 | **0.040** | **0.001** |
| O5 | **0.020** | **0.023** | 0.872 | **0.009** | **0.005** | O5 | **0.005** | **0.032** | 0.401 | 0.367 | **0.001** |
| **c** | -2 °/s | -0.8 | 0 | +0.8 | +2 | **d** | -2 °/s | -0.8 | 0 | +0.8 | +2 |
| All | **0.001** | **0.001** | 0.623 | **0.001** | **0.001** | All | **0.001** | **0.001** | 0.067 | **0.001** | **0.001** |
| O1 | **0.001** | **0.001** | 0.048 | **0.001** | **0.001** | O2 | 0.203 | **0.001** | 0.050 | 0.750 | 0.745 |
| O2 | **0.001** | **0.001** | 0.180 | **0.001** | **0.001** |  |  |  |  |  |  |
| O3 | **0.003** | **0.001** | 0.507 | **0.006** | **0.001** |  |  |  |  |  |  |
| O4 | **0.001** | 0.322 | 0.761 | 0.146 | **0.001** |  |  |  |  |  |  |
| O5 | **0.002** | **0.018** | 0.452 | 0.063 | **0.002** |  |  |  |  |  |  |

**Table S3.** Heading bias for envelope motion <> time-varying phase motion (all comparisons two-tailed). “All”; N observers = 8; N trials per rotation velocity > 960.

|  | -2 °/s | -0.8 | 0 | 0.8 | 2 |
| --- | --- | --- | --- | --- | --- |
| All | 0.523 | **0.009** | 0.157 | 0.948 | 0.537 |
| O2 | 0.359 | 0.418 | 0.493 | 0.107 | 0.274 |

**Table S4.** Heading bias for non-varying (first, initial) > non-varying (first, replication). “All”; N observers = 5; N trials per rotation velocity > 1,200. Each observer “O1-5”, N trials per rotation velocity = 240.

|  | -2 °/s | -0.8 | 0 | 0.8 | 2 |
| --- | --- | --- | --- | --- | --- |
| All | 0.325 | **0.001** | 0.429 | 0.380 | 0.228 |
| O1 | 0.606 | 0.303 | 0.677 | 0.552 | 0.878 |
| O2 | 0.612 | **0.001** | **0.001** | 0.995 | 0.928 |
| O3 | 0.065 | **0.005** | 0.299 | 0.142 | **0.003** |
| O4 | 0.274 | 0.050 | 0.870 | 0.821 | 0.812 |
| O5 | 0.595 | 0.245 | 0.458 | 0.052 | 0.538 |

**Table S5.** Heading bias for non-varying (first) > non-varying (last). **a.** Non-varying (first, initial) > non-varying (last). **b.** Non-varying (first, replication) > non-varying (last). **c.** Non-varying (first, pooled data from **a** and **b**) > non-varying (last). “All”; N observers = 10; N trials per rotation velocity > 1,200.

| **a** | -2 °/s | -0.8 | 0 | +0.8 | +2 | **b** | -2 °/s | -0.8 | 0 | +0.8 | +2 |
| --- | --- | --- | --- | --- | --- | --- | --- | --- | --- | --- | --- |
| All | 0.2537 | 0.4416 | 0.0989 | 0.8921 | **0.001** | All | 0.372 | 1.000 | 0.585 | 0.980 | **0.001** |
| O2 | **0.001** | **0.001** | **0.030** | **0.001** | **0.001** | O2 | **0.001** | 0.999 | **0.001** | **0.001** | **0.001** |

**Table S5 | Heading bias for non-varying (first) > non-varying (last).**
| **c** | -2 °/s | -0.8 | 0 | +0.8 | +2 |
| --- | --- | --- | --- | --- | --- |
| All | 0.237 | 0.970 | 0.270 | 0.931 | **0.001** |
| O2 | **0.001** | 0.332 | 0.960 | **0.001** | **0.001** |

**Movie S1 | Non-varying phase motion (first flow field, 0° heading, no rotation).** Demo video depicting non-varying phase motion, conveying 0° heading (i.e., in the direction of gaze) with 0 °/s rotation (i.e., corresponding to a straight path). For demo purposes, plaid patches have been enlarged ~250% in diameter and reduced ~60% in density. Translation speed, 2.25 m/s, is 1.5x the speed used in experiments.

**Movie S2 | Non-varying phase motion (first flow field, 0° heading, +2 °/s rotation).** Demo video depicting non-varying phase motion, conveying 0° heading (i.e., in the direction of gaze) with 2 °/s rightward rotation (i.e., corresponding to a circular path with radius 43 m). Same modifications as in Movie S1. This convention is used throughout all movies.

**Movie S3 | Non-varying phase motion (last flow field, 0° heading, no rotation).**

**Movie S4 | Non-varying phase motion (last flow field, 0° heading, +2 °/s rotation).**

**Movie S5 | Time-varying phase motion (0° heading, no rotation).**

**Movie S6 | Time-varying phase motion (0° heading, +2 °/s rotation).**

**Movie S7 | Envelope Motion (0° heading, no rotation).**

**Movie S8 | Envelope Motion (0° heading, +2 °/s rotation).**

